# A public-data reproducibility benchmark of a 2.5-D U-Net with a slice-level topology term for type-B aortic dissection segmentation

**DOI:** 10.64898/2026.10.07.757473

**Authors:** Lingsen You, Yujun Guo, Xinyu Zhong, Zisu Peng, Wentong Wang, Li Shen, Junbo Ge

## Abstract

**Background:** Type-B aortic dissection requires separation of the true and false lumina in heterogeneous, anisotropic CTA. A topology term may reduce fragmentation, but its reproducibility must be tested under a fixed case split and independent native-grid evaluation.

**Methods:** We used the public Graz type-B aortic dissection CTA collection (40 cases; CC BY 4.0) and froze a 28/6/6 case split for training, development, and held-out testing. A compact 2.5-D U-Net trained with cross-entropy plus multiclass Dice was compared with the identical model augmented by a fixed-weight (0.5) differentiable slice-level soft clDice term. Two training seeds were run for each arm. All 40 cases passed the NRRD audit; predictions for six held-out cases were restored to their native grids and scored with case-level 3-D voxel-grid Dice, voxel-grid morphological clDice, 6-connected component counts, physical-volume error, and 2,000-resample case-level paired bootstrap intervals.

**Results:** With seed 20261007, topology-minus-baseline differences were −0.028 (95% CI −0.053 to −0.004) for true-lumen Dice, +0.032 (−0.006 to +0.096) for true-lumen voxel-grid clDice, +0.071 (+0.037 to +0.103) for false-lumen Dice, and +0.073 (+0.048 to +0.103) for false-lumen voxel-grid clDice. With seed 20261008, the corresponding differences were −0.040 (−0.065 to −0.018), +0.013 (−0.034 to +0.071), −0.099 (−0.187 to −0.004), and −0.058 (−0.124 to −0.001). Descriptive pooled means over 12 case rows per arm were true-lumen Dice 0.577 for topology versus 0.611 for baseline and false-lumen Dice 0.409 versus 0.422; these pooled rows are not independent cases.

**Conclusions:** Under this fixed six-case test split, the corrected slice-level topology term did not produce a consistent benefit. True-lumen Dice was lower in both seeds, while false-lumen effects reversed direction between seeds; topology-clDice changes were small and/or interval-crossing. The benchmark supports an auditable failure-analysis workflow but does not support a general topology benefit, clinical use, external generalization, or a panvascular decision system. The 3-D clDice quantity is an evaluation descriptor on the voxel grid; the training term is slice-level and does not enforce physical 3-D connectivity.

**Current executed scope and claim boundary:** This executed benchmark is restricted to the 40-case Graz type-B aortic dissection CTA collection with expert true- and false-lumen annotations. All 40 cases passed the audit; the frozen case split contains 28 training, 6 development, and 6 held-out test cases. Four runs (baseline/topology arms with seeds 20261007 and 20261008) produced native-grid predictions for all six held-out cases. The current estimand is descriptive case-level segmentation performance within this single public source. AVT, AortaSeg-60, ImageTBAD, and the ancillary multi-structure CCTA collection are documented as possible future external resources, but they are not part of the current endpoint, training set, or performance claim. This is a proof-of-concept segmentation benchmark, not a panvascular clinical system. The present study does not estimate multicentre generalisation, perform a TEVAR landing-zone assessment, evaluate device fit, or use follow-up outcomes. Any virtual-TEVAR geometry extension remains separately gated future work.

## Introduction

Type-B aortic dissection creates a computational challenge that is clinically relevant and methodologically tractable. A dissection can contain separate true and false lumina, variable thrombus, entry and re-entry tears, and branch-vessel involvement. A model that produces a visually plausible but disconnected false-lumen mask can fail in a way that a global Dice score does not reveal. The first benchmark therefore focuses on the reproducible segmentation problem and keeps any downstream intervention planning outside the present claim boundary.

The device-vessel suitcordance perspective in panvascular intervention treats anatomy, device geometry, imaging measurement, and disease state as a coupled research object [1,6]. The FLOW framework extends panvascular modelling to hemodynamics, longitudinal monitoring, multiple vascular beds, and device interaction [2]. Reviews of intravascular imaging and digital twins likewise emphasize that computational representations must preserve measurement context and validation status [3]. Here those ideas are used to justify explicit spatial provenance and topology checks; the present experiment does not score device-vessel matching and does not perform TEVAR planning.

The AI workflow is also informed by recent scientific-agent research. ScienceBuddy records research interactions, tool calls, execution evidence, and feedback as reusable tasks and evaluation signals. Discovery Foundation Models describe revisable research state, representation construction, evidence-grounded revision, and process-centred evaluation; GALILEO illustrates how such a loop can be grounded in therapeutic-discovery evidence [12–14]. We use these ideas only to structure an auditable procedure loop. We do not reproduce their reinforcement-learning systems, claim recursive self-improvement, or imply that a segmentation model can discover therapies.

The specific question was falsifiable within the Graz cohort: did adding a slice-level clDice objective to an otherwise identical 2.5-D U-Net improve case-level true- and false-lumen topology metrics and volume fidelity over a cross-entropy-plus-multiclass-Dice baseline under a frozen case-level split?

## Objectives and hypotheses

The primary objective was to evaluate true- and false-lumen segmentation on the 40-case Graz collection with a paired, case-level comparison. The secondary objective was to release an auditable data split, grid-validation procedure, and independent metric implementation that can support future external validation. No aortic-tree, thrombus, landing-zone, or clinical-outcome endpoint is part of the current experiment.

The pre-specified hypotheses were:

1. The topology-aware model was hypothesized to improve case-level true- and false-lumen clDice over the standard baseline under the frozen case split; this was tested separately by training seed.
2. Any Dice or volume improvement would be reported separately from topology improvement; a composite would not conceal a class-specific failure.
3. Component counts and physical volume error would identify fragmentation or systematic over- and under-segmentation that a slice average can hide.

## Public data sources and current cohort

### Graz expert-annotated cohort (current benchmark)

The current benchmark uses the public Graz collection of 40 type-B aortic dissection CTA cases with expert true- and false-lumen annotations [8]. The source study searched routine CTAs from a regional hospital network acquired between 2005 and 2021. It excluded type-A dissections, poor image quality, markedly thrombosed false lumina, excessive motion or metal artifacts, scans in which true and false lumina could not be distinguished across more than a few slices, and scans with inadequate contrast. All scans were acquired in the arterial contrast phase, were not ECG-gated, and were reconstructed at 512 x 512 pixels with an average slice thickness of 2.56 mm; 35 of 40 had overlapping slices. The data are anisotropic, so all physical-volume calculations use the NRRD spacing rather than assuming isotropic voxels.

The released files contain anonymized NRRD CT volumes, separate true-lumen and false-lumen NRRD masks, and pathology surface meshes. Entry and re-entry tears are not annotated as labels. Major aortic branches were segmented mainly for orientation and are not a complete branch reference. Three cardiac-surgery residents and one medical student produced the masks using semi-automated 3D Slicer tools followed by substantial slice-wise manual post-processing; student-labelled cases are marked in the source table. These provenance features are retained in the audit manifest and are treated as limitations of the reference standard.

### Future external public cohorts

ImageTBAD (100 cases with true-lumen, false-lumen, and thrombus-related labels), the 56-aorta AVT multicentre aortic-tree cohort, AortaSeg-60, and the 206-case multi-structure CCTA cohort are documented as future external resources [7,9–11]. They are not downloaded into the current experiment, are not used for training or tuning, and are not part of the present performance claim. A future external-validation paper must verify each source’s current access terms, label semantics, and case-level independence before any pooled analysis.

### Access and licence audit

The machine-readable audit records retrieval dates, downloaded-file hashes where available, access routes, licence terms, and known limitations. The current Graz release is CC BY 4.0 and may be downloaded and reused under that licence; the study does not repackage source files. Public availability does not establish clinical representativeness or case-level independence beyond the information supplied by the data owners.

## Preprocessing and case-level splits

Each case is kept as one unit from download through inference. The audit checks CT, true-lumen, and false-lumen size, spacing, origin, and direction before preprocessing. CT and mask geometry were checked and the native direction metadata was retained; the training representation used a fixed full-field-of-view 192 x 192 in-plane resize and three adjacent axial slices as channels. Edge slices use replicated neighbours. No label-derived bounding box, test-time crop, or test-derived intensity statistic is permitted.

The label ontology is background (0), true lumen (1), and false lumen (2). The source masks remain archived unchanged, and any resampling uses nearest-neighbour interpolation for labels. True- and false-lumen overlap is audited before training. For the executable labels, an explicit true-lumen precedence policy was applied to overlapping voxels; overlap counts are retained in the audit and prediction manifests. The audit found 264 overlapping voxels across 16 of 40 cases (maximum 106 in one case).

The public release has no official train/validation/test split. We therefore freeze a case-level split before model training: 28 cases for training, 6 for development, and 6 held-out cases for the primary test. The split file, random seed, case IDs, and the downloaded-file manifest are versioned; source-file hashes are retained in the download manifest. No axial slice, overlapping reconstruction, STL derivative, or mask from a case may cross a split. Preprocessing parameters, augmentation, loss weights, early stopping, and decision thresholds are selected on the training/development cases only. The held-out cases are read once for final inference after checkpoints and analysis code are frozen.

The public files do not provide an independent patient identifier beyond the case directory. The analysis therefore reports case-level results and does not claim verified patient-level independence or that the cases represent 40 unique patients. The split uses case IDs. The public release does not provide a patient-linkage table, so duplicate or near-duplicate patient independence remains unverified; no cross-case merge was inferred from the public directories.

## Models

### Standard 2.5-D baseline

The baseline is a compact 2-D U-Net applied to axial triplets (previous, current, and next slice) with three output classes. The loss is cross-entropy plus multiclass Dice. It uses the same fixed-field-of-view representation, optimizer, batch size, epoch budget, random seeds, and checkpoint rule as the topology arm. The model is a transparent baseline inspired by nnU-Net’s reproducibility principles but is not an nnU-Net implementation [16].

### Slice-level topology arm

The topology arm uses the identical 2.5-D U-Net and training budget, with a fixed 0.5-weight differentiable slice-level soft clDice term added to the cross-entropy plus multiclass Dice loss. For each foreground class, the implementation uses the standard smoothed ratios Tprec = |S(P) ∩ G|/(|S(P)| + ε) and Tsens = |S(G) ∩ P|/(|S(G)| + ε), combined by the smoothed harmonic mean; foreground classes are labels 1 and 2. When a target skeleton is absent, the smoothed topology term is neutral by convention, while cross-entropy and volumetric Dice continue to penalize false-positive probabilities. The term acts on each axial prediction during training; it is not a three-dimensional topology loss and must not be described as 3-D connectivity learning. The clDice weight was fixed a priori at 0.5 for this benchmark and was not tuned on the held-out cases. No hidden centreline or TEVAR label is used.

### Training and checkpoint contract

Prediction manifests record checkpoint and split hashes, native-grid metadata, resize policy, overlap policy, and device/model settings; the exact commands and run configuration are retained in the repository workflow documentation. The executed configuration used 20 epochs, 100 steps per epoch, batch size 16, base width 16, output size 192, AdamW learning rate 2×10-4, and two training seeds. Results in this manuscript come from the complete 40-case audit and the frozen six-case test partition; the smoke run is engineering evidence only.

### Segmentation outcomes

The primary endpoints are case-level Dice and three-dimensional morphological clDice [15] for the true-lumen and false-lumen classes, each reported separately. The evaluator also reports predicted and reference connected-component counts, component-count difference, voxel counts, physical volumes in mm^3^ and ml, signed volume error, absolute volume error, and relative volume error. Both the slice-level training clDice and the 3-D evaluation clDice are labelled explicitly so that they are not conflated.

Dice is defined as (2|P intersect R|/(|P|+|R|)). The evaluation clDice uses 3-D voxel-grid thinning (Lee skeletonization) with topology precision and sensitivity and their harmonic mean. Skeletonization is performed on the voxel grid and is not spacing-aware; this is therefore a morphology descriptor rather than a physically invariant connectivity measure. A class that is absent in both prediction and reference receives a null metric with an explicit ‘both_empty’ status rather than an artificial score of one. A class present in only one mask receives Dice and clDice equal to zero. All connected-component and volume metrics are calculated on the native prediction/reference grid after grid compatibility checks. Component counts use 6-connectivity; physical volumes use the NRRD voxel spacing.

A component count is not a clinical endpoint. It is a failure descriptor that makes fragmentation visible even when voxel overlap is acceptable. Volume error is a measurement error relative to the supplied reference mask and does not establish an anatomic or treatment threshold.

## Results

### Data audit and execution

The canonical Figshare record established the provenance and CC BY 4.0 terms; the case files were retrieved from the public mirror listed in the manifest, and all 40 case directories passed the executable NRRD audit (40 discovered, 40 valid, 0 errors, 40 cached). True/false-lumen overlap was detected in 16 cases (264 voxels in total; maximum 106 voxels in one case) and resolved for this run by the predeclared true-lumen-precedence policy. The frozen split contained 28 training, 6 development, and 6 held-out cases. All four model/seed arms completed training and produced valid native-grid predictions for each of the six held-out cases: cta01s, cta05s, cta17s, cta28s, cta33s, and cta39s. No prediction was imputed or excluded for grid mismatch.

### Held-out performance

The independent evaluator summarized the six held-out cases for each arm. Values are means across cases; the two seeds are shown separately because they are repeated training runs, not additional independent cases.

| arm | training seed | TL Dice | TL voxel-grid cIDice | FL Dice | FL voxel-grid cIDice | FL predicted components |
| --- | --- | --- | --- | --- | --- | --- |
| baseline | 20261007 | 0.583 | 0.405 | 0.381 | 0.261 | 262.0 |
| slice-level topology | 20261007 | 0.554 | 0.437 | 0.452 | 0.335 | 90.8 |
| baseline | 20261008 | 0.639 | 0.515 | 0.464 | 0.387 | 152.2 |
| slice-level topology | 20261008 | 0.599 | 0.528 | 0.366 | 0.328 | 42.7 |

The topology arm reduced predicted false-lumen component counts in both seeds (90.8 versus 262.0 in seed 20261007; 42.7 versus 152.2 in seed 20261008), but this descriptor did not track a stable anatomy benefit: false-lumen Dice and clDice improved in seed 20261007 and declined in seed 20261008. Relative volume errors were +25.5% versus +34.6% (topology versus baseline) in seed 20261007 and −47.1% versus −26.8% in seed 20261008. Component count alone therefore cannot be interpreted as improved segmentation. Across both seeds pooled descriptively (12 case rows per arm), true-lumen Dice was 0.577 for topology versus 0.611 for baseline, whereas false-lumen Dice was 0.409 versus 0.422. These pooled values are descriptive only and are not a confirmatory estimate.

### Case-level paired effects

A 2,000-resample paired bootstrap was run separately for each training seed on the six common held-out cases. The effect is topology minus baseline; no p-values or multiplicity-adjusted claims are reported.

| seed | metric | mean difference | percentile 95% CI |
| --- | --- | --- | --- |
| 20261007 | TL Dice | -0.028 | -0.053 to -0.004 |
| 20261007 | TL voxel-grid cIDice | +0.032 | -0.006 to +0.096 |
| 20261007 | FL Dice | +0.071 | +0.037 to +0.103 |
| 20261007 | FL voxel-grid cIDice | +0.073 | +0.048 to +0.103 |
| 20261008 | TL Dice | -0.040 | -0.065 to -0.018 |
| 20261008 | TL voxel-grid cIDice | +0.013 | -0.034 to +0.071 |
| 20261008 | FL Dice | -0.099 | -0.187 to -0.004 |
| 20261008 | FL voxel-grid cIDice | -0.058 | -0.124 to -0.001 |

The direction of the topology effect changed across seeds for both false-lumen metrics, while true-lumen Dice decreased in both seeds and topology-clDice changes were small. This seed sensitivity is the principal result and prevents a general claim that the slice-level topology term improves type-B lumen segmentation. The initial pre-correction run was excluded after the clDice ratio and soft-skeleton implementation audit; only the corrected outputs are reported here.

## Statistical analysis

The primary comparison was the case-level paired difference between the topology arm and the baseline on the four class-metric pairs (true-lumen Dice, true-lumen voxel-grid clDice, false-lumen Dice, and false-lumen voxel-grid clDice). Secondary descriptors were component counts and physical volume errors. The evaluator retained every case, class, and status; all 24 arm-by-case-class evaluations for each seed were valid, with no missing predictions or grid mismatches.

For each metric and class, 2,000 case-level paired bootstrap resamples were drawn with replacement from the six eligible held-out cases. The reported effect is topology minus baseline with a percentile 95% interval. The bootstrap random-number seed was fixed at 20261007 for reproducibility and is distinct from the two training seeds. Slices were never bootstrap units. No p-value, multiplicity adjustment, or composite score was introduced after observing the test results. Because the public release supplies case directories rather than a verified independent patient identifier, the analysis is case-level and does not claim patient-level independence. The six-case estimates are therefore a precision-limited reproducibility benchmark rather than evidence of clinical efficacy or broad generalization.

## Feedback-aware procedure loop

The procedure loop is limited to reproducibility and human-supervised audit. It contains a fixed model, an execution record, a verifier, and a reviewer-visible change log. A proposed change can affect preprocessing, loss weighting, or checkpoint selection, but it may be accepted only on the training/development cases with the held-out test partition untouched. Every accepted and rejected candidate, its parent version, and its evidence are retained. This workflow is informed by the auditable research-state ideas of ScienceBuddy and Discovery Foundation Models [12,13]; it does not claim recursive self-improvement, reinforcement learning, or autonomous discovery.

## Relation to the panvascular AI programme

PanVasc-Suit narrows the device-vessel suitcordance idea to an auditable imaging task. A literature or registry agent must preserve evidence provenance, whereas a segmentation agent must preserve spatial topology, native-grid metadata, and measurement uncertainty. The two tasks may eventually share a research state containing source, image, segmentation, geometry, protocol, and reviewer feedback, but this Graz benchmark does not claim that such an integrated system has been built.

The Biomarker Research cohort paper is cited as a reminder that clinical analyses must preserve treatment allocation, observed outcome, and adjustment strategy [5]. It is not used as a segmentation label or as evidence for TEVAR benefit. The panvascular scaffold-design study is similarly cited as computational context, not evidence for the segmentation endpoint [4]. Any geometry or device-suitcordance analysis is future work after an independent external dataset and a separately approved analysis plan.

## Figures and reproducibility artifacts

The executed benchmark generated three figures from the frozen manifests and run outputs:

**Figure 1.**
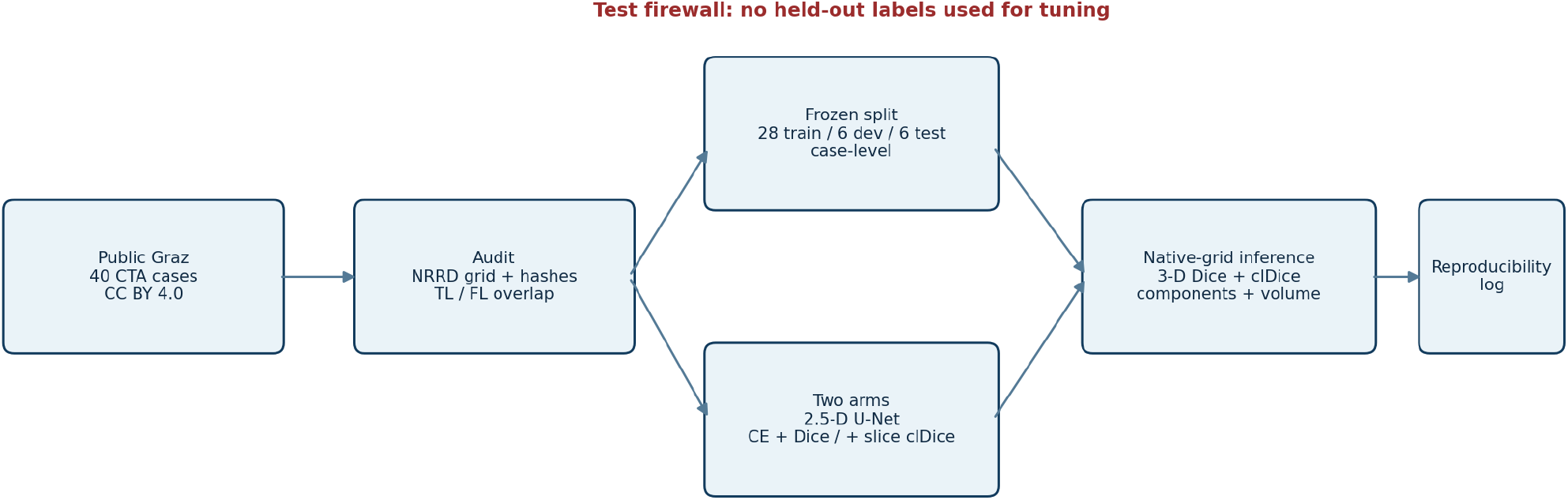
Study flow and audit boundary. The 40-case Graz retrieval, licence check, native-grid validation, case-level split, two model arms, held-out inference, independent evaluation, and reproducibility checkpoints are shown.

**Figure 2.**
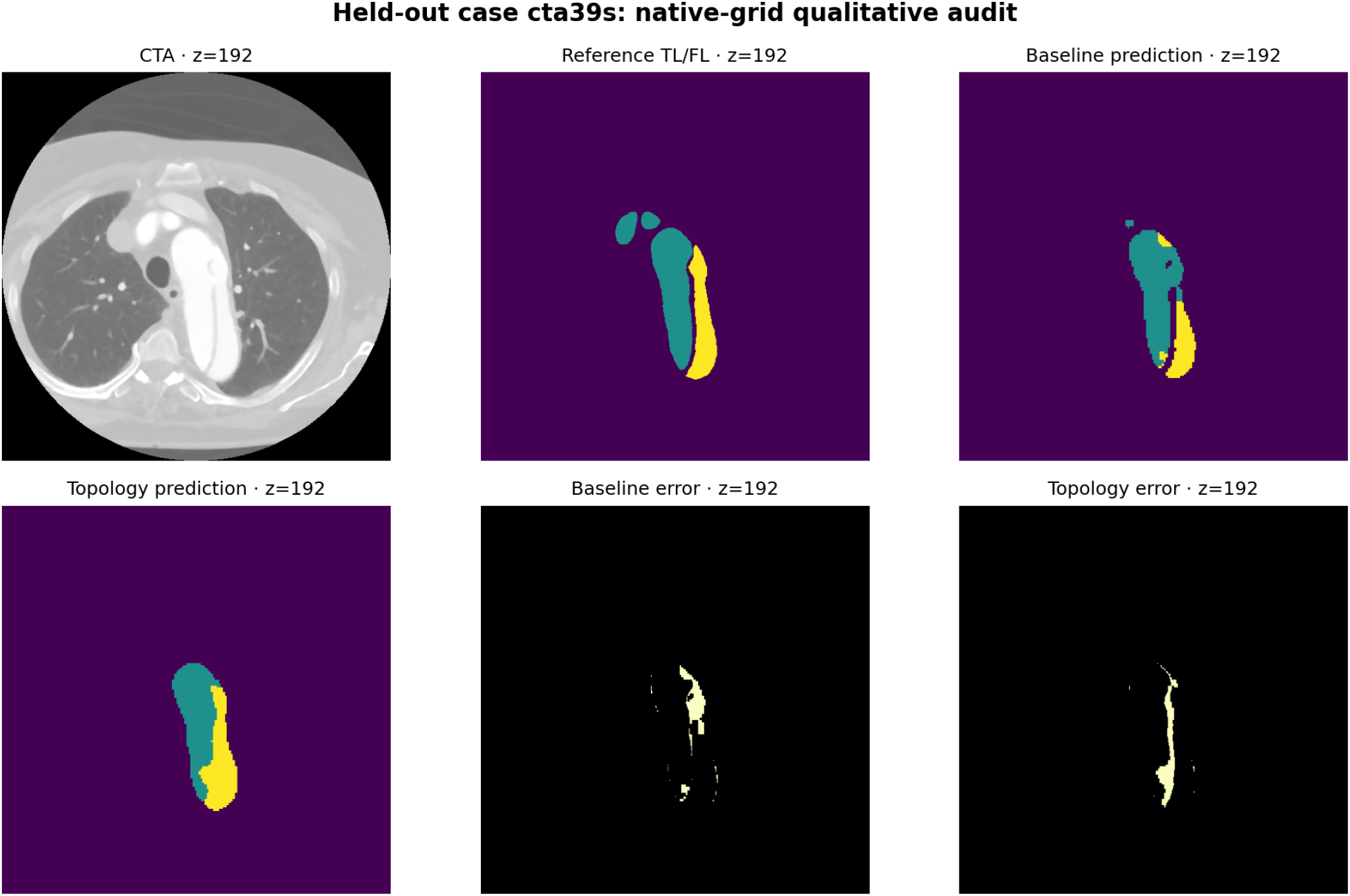
Representative held-out segmentation. Axial CTA, reference TL/FL masks, baseline prediction, topology prediction, and error panels for held-out case cta39s are shown. This is an illustration of the failure-analysis workflow, not a substitute for case-level summaries.

**Figure 3.**
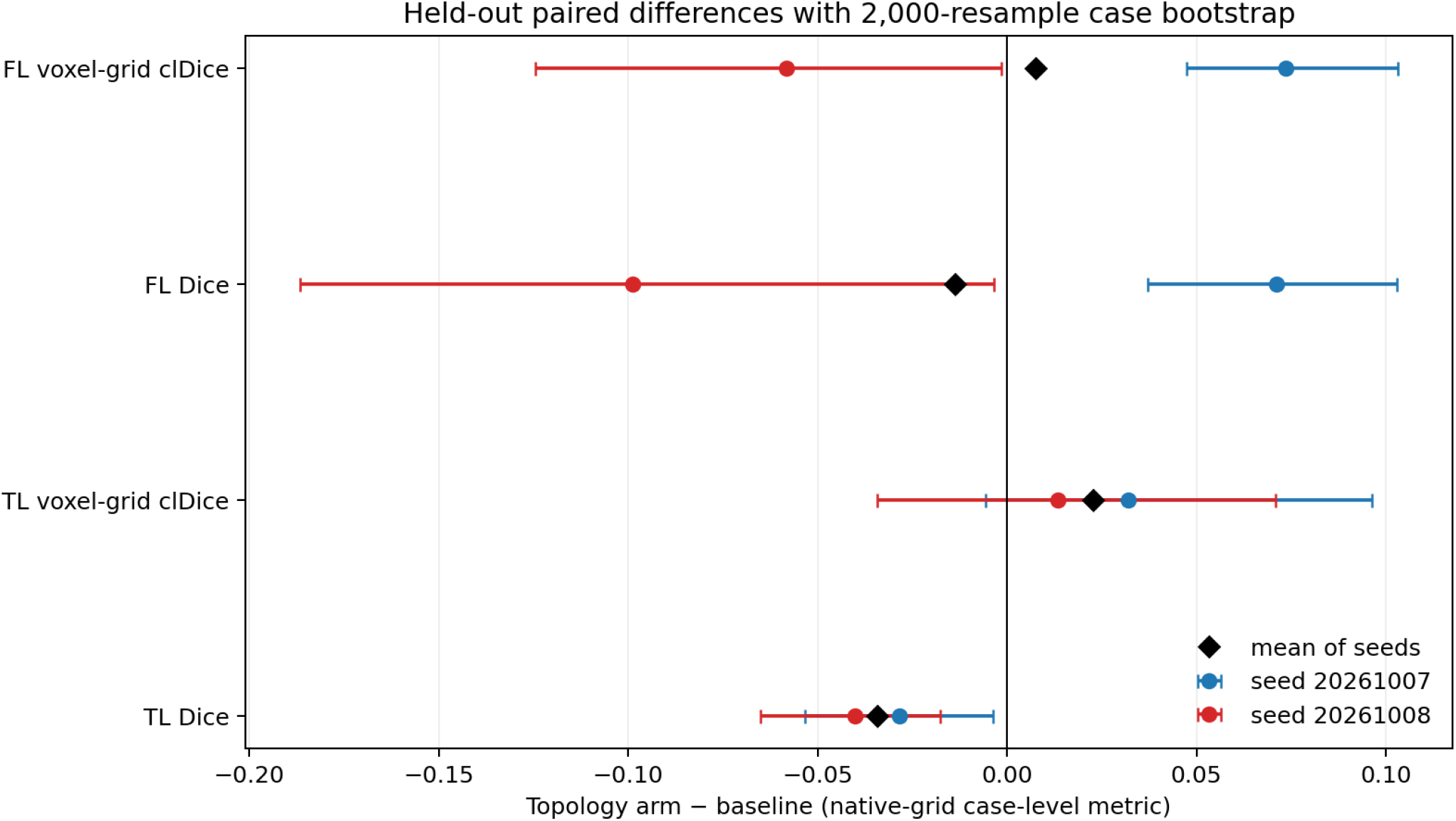
Paired case-level comparison. Seed-specific topology-minus-baseline effects with 95% paired bootstrap intervals are shown.

The repository provides results_corrected_v3/case_metrics.csv, results_corrected_v3/summary.json, the three figures, the fixed split, data and checkpoint manifests, the native-grid evaluator, and the bootstrap script. No figure displays a TEVAR device, landing-zone recommendation, or simulated result. The legacy results/ directory contains pre-correction outputs and is excluded from the manuscript. The output directory is an analysis artifact; the public source images and masks are not redistributed.

## Limitations and safety boundary

This is a small, single-source public cohort selected for clear TL/FL delineation. The source study excluded many difficult scans, so the benchmark cannot estimate performance on all type-B dissections or on cases with severe thrombosis, poor contrast, metal artifacts, or ambiguous lumen identity. The released files do not establish independent patient IDs beyond case directories, and the source annotation process involved semi-automated tools and substantial manual editing by multiple annotators. The masks do not provide complete entry/re-entry, thrombus, or branch reference labels.

The 2.5-D model sees neighbouring axial slices but does not enforce three-dimensional topology during training, and the voxel-grid clDice evaluator is not spacing-aware. The true-lumen-precedence overlap policy may affect a small number of reference voxels and is not an adjudicated clinical correction. The topology arm was seed-sensitive, with opposite false-lumen effects across the two seeds despite fewer predicted components in both topology arms. A six-case held-out partition yields wide uncertainty and is suitable for a reproducibility benchmark, not a clinical validation. No external cohort, multicentre generalisation, thrombus, entry/re-entry tear, branch, device-fit analysis, TEVAR landing-zone calculation, follow-up outcome, safety, efficacy, or patient-benefit claim is made. Any such analysis requires a new protocol, independent data, and clinical review.

## Data and code availability

The working repository includes the Graz accession manifest, licence and access decisions, case-level split files, preprocessing scripts, 2.5-D model configurations, native-grid validation code, evaluate_predictions.py, paired_bootstrap.py, aggregate outputs, figure-generation code, and the downloaded-file manifest with hashes. Restricted files will not be redistributed. Credentials, patient identifiers, and private clinical images are excluded. The released evaluator retains missing-case and grid-mismatch records rather than silently filtering them.

## Funding, competing interests, and AI use

This project was supported by the National Natural Science Foundation of China (T2288101, 82170342) and the Medical Engineering Joint Fund of Fudan University (yg2023-01). The language of this article has been polished with the assistance of AI-based language tools.

Lingsen You, Li Shen, and Junbo Ge have participated in development of the XINSORB bioresorbable scaffold. This study evaluates no XINSORB device or other proprietary implant. The remaining authors declare no competing interests.

## DECLARATION OF GENERATIVE AI AND AI-ASSISTED TECHNOLOGIES

Codex assisted with LaTeX assembly, prose editing, and consistency checks. The authors reviewed and revised all AI-assisted text and take responsibility for the submitted manuscript.

## Author contributions

Lingsen You: Conceptualization, Methodology, Software, Formal analysis, Visualization, Writing - original draft. Yujun Guo: Methodology, Data curation, Validation, Writing - original draft. Xinyu Zhong: Data curation, Methodology, Validation, Writing - original draft. Zisu Peng: Data curation, Formal analysis, Visualization, Writing - original draft. Wentong Wang: Data curation, Validation, Writing - review and editing. Li Shen: Conceptualization, Supervision, Funding acquisition, Writing - review and editing. Junbo Ge: Conceptualization, Supervision, Funding acquisition, Project administration, Writing - review and editing. All authors reviewed and approved the final manuscript and are accountable for the work.

